# Visual deprivation during mouse critical period reorganizes network-level functional connectivity

**DOI:** 10.1101/2023.05.30.542957

**Authors:** Siyu Chen, Rachel M. Rahn, Annie R. Bice, Seana H. Bice, Jonah A. Padawer-Curry, Keith B. Hengen, Joseph D. Dougherty, Joseph P. Culver

**Affiliations:** Department of Radiology, Washington University School of Medicine, St. Louis, MO 63110, USA; Department of Genetics, Washington University School of Medicine, St. Louis, MO 63110, USA; Department of Psychiatry, Washington University School of Medicine, St. Louis, MO 63110, USA; Department of Biology, Washington University, St. Louis, MO 63130, USA; Intellectual and Developmental Disabilities Research Center, Washington University, St. Louis, MO 63130, USA; Department of Physics, Washington University School of Medicine, St. Louis, MO 63110, USA; Department of Biomedical Engineering, Washington University School of Medicine, St. Louis, MO 63110, USA

**Keywords:** Monocular deprivation, functional connectivity, calcium neuroimaging, plasticity, cortical reorganization.

## Abstract

A classic example of experience-dependent plasticity is ocular dominance (OD) shift, in which the responsiveness of neurons in the visual cortex is profoundly altered following monocular deprivation (MD). It has been postulated that OD shifts also modify global neural networks, but such effects have never been demonstrated. Here, we used longitudinal wide-field optical calcium imaging to measure resting-state functional connectivity during acute (3-day) MD in mice. First, delta GCaMP6 power in the deprived visual cortex decreased, suggesting that excitatory activity was reduced in the region. In parallel, interhemispheric visual homotopic functional connectivity was rapidly reduced by the disruption of visual drive through MD and was sustained significantly below baseline state. This reduction of visual homotopic connectivity was accompanied by a reduction in parietal and motor homotopic connectivity. Finally, we observed enhanced internetwork connectivity between visual and parietal cortex that peaked at MD2. Together, these findings support the hypothesis that early MD induces dynamic reorganization of disparate functional networks including association cortices.

**Significance Statement:** Monocular deprivation during the visual critical period triggers several plasticity mechanisms that collaborate to shift the excitability of neurons in the visual cortex. However, little is known about the impacts of MD on cortex-wide functional networks. Here, we measured cortical functional connectivity during short-term critical period MD. We demonstrate that critical period MD has immediate effects on functional networks beyond the visual cortex, and identify regions of substantial functional connectivity reorganization in response to MD.

## Introduction

Decades of research have revealed that several dramatic forms of plasticity occur in the visual cortex following monocular deprivation (MD) during early postnatal development. In rodents, MD-induced ocular dominance (OD) shifts follow a biphasic time course, where the early phase suppresses deprived-eye responses, and the later phase strengthens both the deprived and open-eye responses (Cooke & Bear, 2014; Frenkel & Bear, 2004; Mrsic-Flogel et al., 2007). It is well-established that the initial suppression of deprived-eye responses results from long-term depression (LTD)-like mechanisms (Heynen et al., 2003; Yoon et al., 2009), as well as plasticity mechanisms in inhibitory circuits (Keck et al., 2011; Maffei et al., 2006, 2010). Recently, it has also been shown that callosal inhibition of deprived-eye responses contribute to OD shifts during MD (Pietrasanta et al., 2014; Restani et al., 2009). However, the possibility that this manipulation may impact disparate functional networks remains a critical and untested prediction of neural plasticity. Here, we used wide-field optical imaging (WFOI), combined with genetically encoded calcium indicators (GECIs), to test whether acute MD and its subsequent plasticity processes cause reorganization of resting-state functional connectivity (rsFC) in the neocortex.

rsFC – the temporal correlation of spontaneous activity between different brain regions – reflects the organization of functional networks. In humans, resting-state functional magnetic resonance imaging (rs-fMRI) studies have shown that changes in rsFC patterns exhibit feed-forward mechanisms and thus reflect prior experience (Grigg & Grady, 2010; Hasson et al., 2009; Waites et al., 2004). For example, sensory deprivation due to upper limb casting (Newbold et al., 2020), congenital and early blindness (Burton et al., 2014; Wang et al., 2013), intensive training on relational reasoning (Mackey et al., 2013), and visual perceptual learning (Lewis et al., 2009) have all been shown to alter rsFC organization. However, there has been limited characterization of rsFC changes during OD plasticity. *In vivo* anesthetized optical intrinsic signal imaging after 14-day MD found a decrease in interhemispheric visual homotopic connectivity but small changes in distant networks (Kraft et al., 2017). However, measuring end-point rsFC using hemodynamic signals from anesthetized animals may not be the best probe for network-level consequences for a number of reasons. Most critically, because Hebbian and homeostatic mechanisms operate with distinct temporal profiles (Hengen et al., 2013, 2016; Rittenhouse et al., 1999), sampling only once after 14 days MD may miss most of the time course of network re-programming. In addition, blood-based intrinsic signal imaging operates on the assumption that hemodynamics can be used as a surrogate reporter of neuronal activity, which may become less reliable during early postnatal periods when neurovascular coupling may still be developing. Therefore, we set out to perform calcium-based WFOI to map the differential responses in rsFC during the early phase of MD. We hypothesized that MD-induced OD shifts would have circuit-level impacts on distant networks, contrasting this with the alternative hypothesis that the loss of visual drive would only modify the immediate circuit.

To test these hypotheses, we took advantage of a classic MD paradigm using lid suturing to perturb visual experience in juvenile mice during visual critical period (postnatal day 24-27 [P24-P27]), a developmental window where MD is known to induce different forms of plasticity (Gordon & Stryker, 1996; Mrsic-Flogel et al., 2007; Smith et al., 2009; Yoon et al., 2009). To avoid the confounds of anesthesia, we performed awake resting-state calcium imaging using transgenic *Thy1*-GCaMP6f mice. We performed imaging before lid suturing and then repeated imaging for three days thereafter, and replicated our findings using both within-subjects (MD vs. baseline) and between-subjects (sham vs. suture) analyses. Overall, we found changes in both visual within-network relationships and distinct between-network relationships. These results support the hypothesis that neuronal changes during OD shifts have large-scale network-level impacts that alter resting-state functional connectivity, and provide a description of the network-wide consequences of attenuated visual input during a visual critical period.

## Materials and Methods

### Animals

All procedures described below were approved by the Washington University Institutional Animal Care and Use Committee and conducted in accordance with the approved Animal Studies Protocol. Breeding animals were kept in a facility with a 12:12 light/dark cycle, in standard cages with nestlets and a cardboard Bio-Tunnel (Bio-Serv) and *ad libitum* access to standard lab diet and water. Adult *Thy1*-GCaMP6f (JAX:024276) homozygous males and females were bred to heterozygous *Myt1l* knockout females and males without the GCaMP6f allele to produce animals that were hemizygous for the *Thy1*-GCaMP6f allele to enable calcium imaging. Genotyping for *Myt1l* was used to exclude mutant animals from analysis such that only wildtype pups were used for analyses here.

Experimental animals were weaned at postnatal-day 21(P21) and housed by sex. All experimental animals were kept in standard cages with *ad libitum* access to food and water and enrichment, including a mouse tube (Bio-Serv), mouse hut (Braintree Scientific), nestlets and hydrogel (Bio-Serv).

### Cranial windowing

A transparent chronic optical window made of plexiglass was fitted to the dorsal cranium of the mouse at P21 as previously described (Rahn et al., 2021). Briefly, the mouse was anesthetized via isoflurane and the head was shaved. An incision was then made along the midline of the scalp to retract the skin and expose an approximately 1.1 cm^2^ dorsal cortical field of view. The plexiglass window was adhered to the dorsal cranium using Metabond clear dental cement (C&B-Metabond, Parkell Inc., Edgewood, NY). Mice were returned to their cages to recover for at least 36 hours before they were habituated head-fixed in the imaging apparatus at P23.

### Eyelid suturing

Lid suturing was performed between zeitgeber time (ZT) 10-15, after baseline imaging on P24. Prior to lid suturing, mice were anesthetized using isoflurane and received 5mg/kg of carprofen (RIMADYL®) for analgesia via subcutaneous injection. Eyelashes and lid margins were first trimmed using Vannas scissors. 3-4 mattress sutures were then applied using sterile 6-0 prolene sutures (Ethicon). Sham MD mice were subjected to the same length of anesthesia and received carprofen injection. After suturing, mice were returned to clean cages with cagemates and enrichment, and 0.5 mg carprofen tablet (Bio-Serv) was placed on the cage floor for each mouse. All mice continued to receive 0.5 mg carprofen tablet for three days after lid suturing surgery. Suture integrity was checked prior to each imaging session. Mice who developed eye infections or whose sutures opened prematurely were dropped from the experiment.

### Wild-field fluorescence optical imaging

Mice were imaged in a wide-field optical imaging system with four sequential illumination via LEDs (470 nm, 530 nm, 590 nm, and 625 nm) which allowed for measuring calcium fluorescence as well as hemodynamic fluctuations (Wright et al., 2017). An sCMOS camera (Zyla 5.5, Andor Technologies; Belfast, Northern Ireland, United Kingdom) captured frames at a rate of 16.8Hz per LED channel, which allowed for analysis in the delta frequency band (0.4-4Hz). The first two cohorts of mice had five 5-minute imaging runs of resting-state awake data collected, while all other cohorts had four 5-minute imaging runs collected to better manage imaging workload. All data collection was done between ZT2 and ZT10. To reduce stress and movement artifacts, all mice were acclimated to head-fixed imaging at P23 using the experimental protocol.

### Imaging data processing

Data were processed using the previously published optical imaging toolbox in MATLAB (Brier & Culver, 2023). Briefly, a binary brain mouse was created by manually tracing the mouse dorsal cortex within the field of view. Raw data were spatially downsampled to 78×78 pixels^2^ and temporally downsampled by 2, and a representative frame of background light levels was subtracted from the data. Each pixel’s individual time trace was then temporally and spatially detrended. Using the changes in the reflectance in the 530 nm, 590 nm and 625 nm LED channels, a modified Beer-Lambert Law was solved to generate fluctuations in oxy- and deoxy-hemoglobin concentration, as described previously (White et al., 2011). Hemodynamic absorption of GCaMP6 emission was corrected using a previously published method (Ma et al., 2016). Spatial smoothing was done with a Guassian filter (5.5 pixel box with a 1.2 pixel standard deviation)., and global signal regression was performed by regressing the average of all time traces of pixels within the mask from each pixel’s time trace. Images were then affine transformed to a common atlas place based on the Paxinos atlas using the locations of bregma and lambda. All analyses were performed on data bandpass filtered to the 0.4-4 Hz delta band using a Butterworth filter.

To exclude motion artifacts, individual 5-minute runs were excluded from analysis if raw LED light levels showed greater than 1% variance across the run, which suggests high levels of movement. Following this motion censoring, 5-25 minutes of data from each mouse were used for analysis at each timepoint.

### Functional connectivity

Three measures of functional connectivity were computed. Pixel-based homotopic interhemispheric rsFC was evaluated by calculating the correlation coefficient between each pixel’s individual time trace and its contralateral pixel’s time trace (i.e. homotopic connectivity). Seed-based rsFC was calculated using a set of 26 cortical seeds as described previously (Brier et al., 2023; Rahn et al., 2021). Seed-based rsFC maps were constructed by calculating the Pearson correlation coefficients between the time trace of a seed region and that of other pixels in the brain mask. Lastly, a matrix approach was taken to examine changes in the functional connections between a total of 325 seed pairs by calculating the Pearson correlation coefficients between the time traces within two seed regions.

### Power spectral analysis

To determine the amount of spontaneous GCaMP6 activity in the delta frequency band (0.4-4Hz), power spectral analysis was performed by splitting delta GCaMP6 time-series at each pixel into 10s segments within each 5-min run. A Hann window was applied to those 10s segments and Fast Fourier Transform (FFT) calculation was performed. The final power spectra was obtained by squaring the FFT.

### Statistical Analysis

A repeated measures ANOVA (rmANOVA) and paired *t*-tests were performed for within-subject analyses at each MD timepoint versus baseline, and thus only mice with data collected at all time points were included in the analysis. A two-way rmANOVA was used to compare changes in correlations over time for MD versus sham groups. Two-sample *t*-tests were used to compare MD mice to sham mice at each timepoint. For two-dimensional maps, a cluster size-based thresholding method adapted from (Brier & Culver, 2023) was used to correct for multiple comparisons and determine statistical significance. This cluster size-based approach credits larger clusters with greater statistical significance than smaller clusters that have the same peak *t* values.

## Results

### Mapping rsFC with Calcium-Based WFOI

We characterized resting-state functional networks of pyramidal neurons during OD shifts using the *Thy1-*GCaMP6f mouse line. We first performed baseline imaging on postnatal day 24 (P24) in order to establish rsFC in the unperturbed brain **(Fig. 1A)**. We then manipulated visual experience via right lid suturing on late P24, and sutures were maintained for 3 days (through P27) as OD shifts are reliably produced during this developmental window (Smith et al., 2009). We repeated optical imaging at the following time points: 18-22 hr monocular deprivation day 1 [MD1]), 42-46 hr (MD2), and 66-70 hr (MD3) post lid suturing (n=20). A schematic of the experimental design is depicted in **Fig. 1A**. As an additional control, another cohort of mice (n=15) underwent sham suturing after baseline imaging. Throughout this study, our analyses thus implemented a within-subject design by comparing each mouse to its own baseline. For completeness, we compared the changes from baseline in MD and sham-sutured mice to enable us to better interpret results and distinguish MD-mediated effects from any natural developmental changes in this time window. Overall, we collected 140 sessions of imaging data from 35 mice in total, representing 40 total hours of data for subsequent analysis.

**Figure 1.**
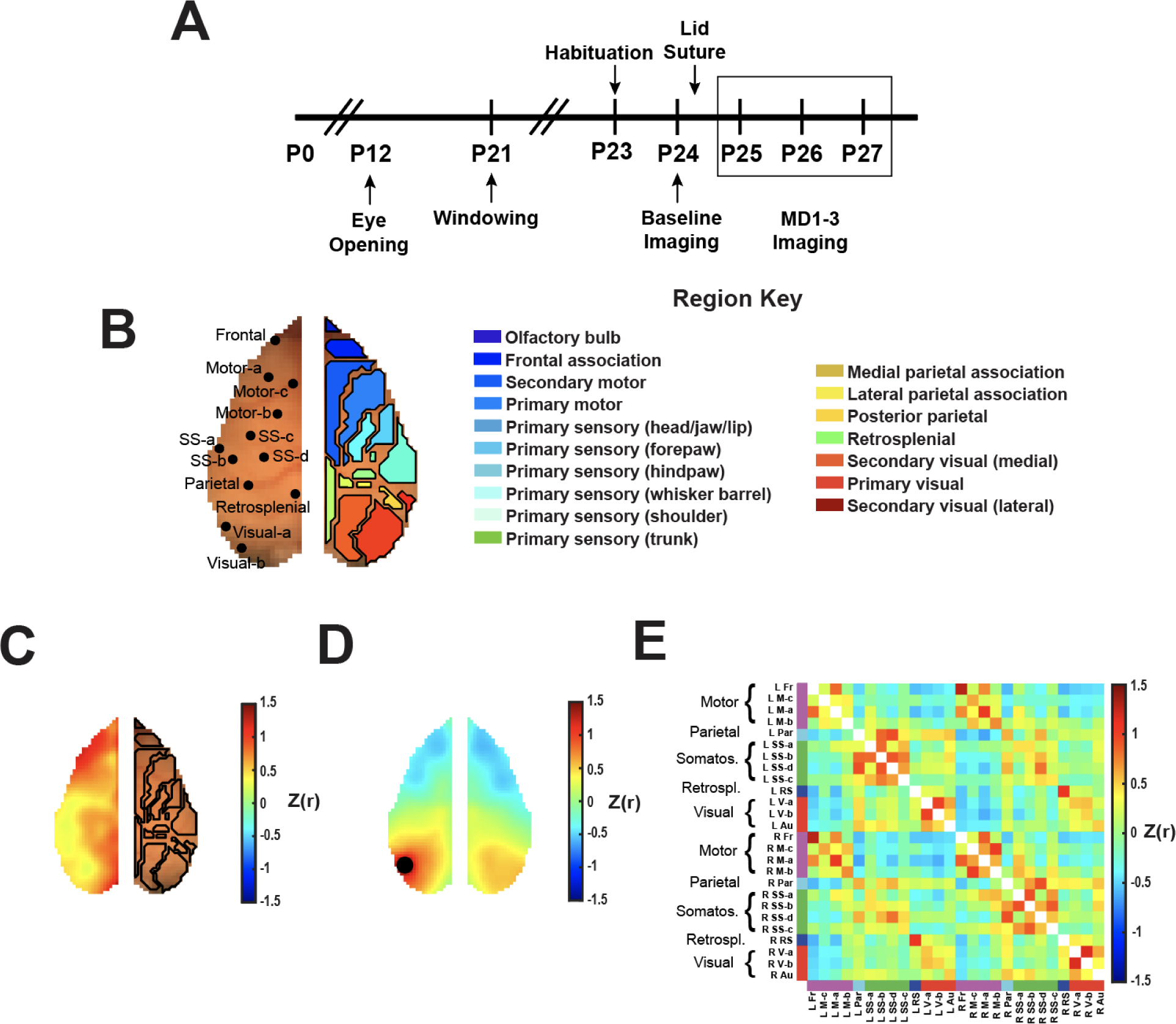
Monocular deprivation (MD) and calcium-based WFOI paradigm. **(A)** Experimental timeline for optical imaging and lid suturing. For MD mice (n=20;14 females and 6 males), the right eyelid was sutured after baseline imaging at P24. Sham-sutured mice (n=15; 5 females and 10 males) were subjected to the same length of isoflurane anesthesia. Mice were subsequently imaged at MD1 (P25), MD2 (P26), and MD3 (P27). **(B)** Visualization of sampled seed centers overlaid with Paxinos atlas brain regions. **(C)** Representative baseline interhemispheric homotopic connectivity map derived from sample 5-min run data, displaying Fisher z-transformed Pearson’s correlation coefficients between each pixel in the left hemisphere and its corresponding pixel in the right hemisphere. Only the left hemisphere is displayed, as homotopic connectivity is symmetrical across the midline. **(D)** Representative baseline left visual seeded rsFC map derived from 5-min run data. Black dot represents the Visual-a seed region, and colormap represents correlation to that seed of each pixel’s time traces. **(E)** rsFC matrix from the same 5-min run data displaying correlation values for each seed pair.

To understand how rsFC reorganizes following sensory deprivation, we investigated changes to four complementary measures of rsFC: pixel-based interhemispheric homotopic functional connectivity maps **(Fig. 1C)**, seed-based rsFC maps **(Fig. 1D)**, and seed-based correlation matrices **(Fig. 1E)**. Interhemispheric homotopic rsFC reflects cortex-wide temporal correlation between homotopic contralateral pixel pairs, and could illustrate changes between the ipsilateral and contralateral visual cortex. Seed-based rsFC refers to the temporal correlation between the seeded region of interest (ROI) and all other pixels in the field of view (FOV), here allowing a focused examination of the deprived visual cortex’s connections to the rest of the brain. Node-degree quantifies rsFC strength, representing the number of positive connections each pixel exhibits with other areas in the cortex. Finally, the correlation matrix examines the correlation between 26 canonical seeds thereby reflecting global changes in rsFC after MD, even in connections not involving visual seeds. If the primary hypothesis is correct, then we predict that MD would result in global rearrangements that manifest as changes in all four measures of FC. If the alternative hypothesis is correct, then the consequence of the loss of input would be restricted to changes within the visual cortices.

### MD Reduces Delta GCaMP6 Power in the Deprived Visual Cortex

To confirm that MD reduces spontaneous neuronal activity as expected, we examined delta GCaMP6 power, as it was previously reported that short-term critical period MD decreases delta in the visual cortex when measured by local field potentials (Malik et al., 2022). We therefore investigated the changes in cortex-wide delta GCaMP6 power in both MD and sham mice and compared the effects between the two groups. To determine the effect of MD versus natural development on delta GCaMP6 power, we first plotted power as a percent of baseline at each MD timepoint for both sham and MD mice **(Fig. 2B & 2C)**. In MD mice, the reduction of delta GCaMP6 power was restricted to the left (deprived) visual cortex **(Fig. 2B)**. In contrast, sham mice displayed an increase of delta GCaMP6 power in both visual hemi-cortices **(Fig. 2C),** suggesting this is an aspect of development. Thus, next, to distinguish the effect of MD from the effect of development, we compared the changes in delta GCaMP6 power between MD and sham mice **(Fig. 2D)**. We found that MD significantly reduced delta GCaMP6 power in the deprived visual cortex at MD3 **(Fig. 2E)**, confirming an expected reduction in spontaneous GCaMP6 activity due to MD. To investigate the changes in connectivity of the deprived visual cortex where activity was reduced, we placed a seed in the cluster of pixels with *t* < −2 for subsequent visual-seeded rsFC analysis **(Fig. 2F)**.

**Figure 2.**
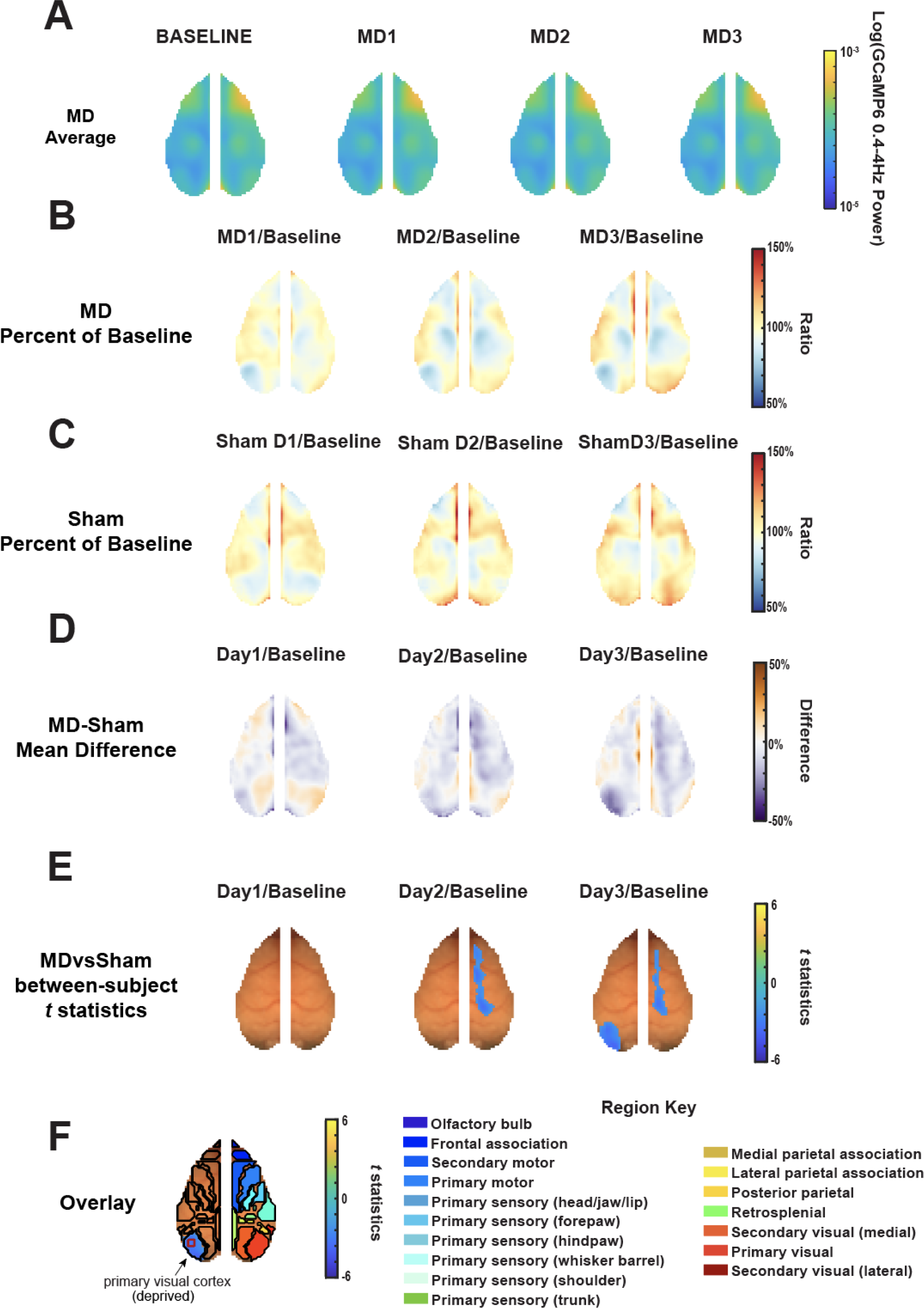
3-Day Critical period monocular deprivation (MD) decreases delta frequency GCaMP6 power in the visual cortex. **(A)** Average MD delta GCaMP6 power maps at each timepoint (n=20). **(B)** MD delta GCaMP6 power plotted as the averaged percentage of each mouse’s baseline values. **(C)** Sham delta GCaMP6 power plotted as the averaged percentage of each mouse’s baseline values (n=15). **(D)** Mean difference maps comparing the changes in delta GCaMP6 power from baseline between MD and sham mice that were shown in (B) and (C). **(E)** Two sample *t*-tests between each MD time point and baseline. Maps display pixels that had significant change in delta GCaMP6 power. **(F)** Overlay of the significant cluster in the visual cortex with Paxinos-atlas based region outlines. Red square displays the visual seed for subsequent analysis.

### MD Suppresses Visual Homotopic Functional Connectivity

The early phase of MD is known to produce a strong and rapid shift in responsiveness of visual cortex neurons in favor of the open eye (Mioche & Singer, 1989). This shift toward the open eye is attributable to the induction of LTD in the visual cortex (Heynen et al., 2003; Liu et al., 2008; Yoon et al., 2009) as well as callosal suppression of deprived-eye responses (Restani et al., 2009). We therefore hypothesized that MD would cause a rapid decrease in visual homotopic connectivity, as asymmetric visual experience would cause spontaneous neuronal activities from the two hemispheres to become desynchronized. However, over time if responsiveness of the deprived visual cortex neurons returns to the contralateral eye, homotopic connectivity should also return.

We were interested specifically in the temporal response of homotopic connectivity to the manipulation of visual experience, and so we investigated the changes in homotopic contralateral rsFC relative to each animal’s baseline at each timepoint **(Fig. 3A, 3B & 3C)**. Because prior work has shown rsFC naturally changes across this time period in rodents (Rahn et al., 2021), we then compared the differences in those changes between MD and sham groups using a two-sample *t*-test **(Fig. 3D & 3E)**. We found that homotopic connectivity between the deprived and non-deprived visual cortices displayed a significant decrease in MD mice compared to sham mice at MD2 and MD3 **(Fig. 3E)**. In addition, pixels in parietal and sensory regions, as well as those in frontal and motor regions, showed a decrease in homotopic connectivity during MD compared to sham mice **(Fig. 3E)**. We examined the correlations between the contralateral visual seed pairs using a two-way rmANOVA **(Fig. 3F & 3G**, **Table 1)**. Both of our seed pairs in the visual cortices (Visual-a and -b seeds, see **Fig 1B** for key) exhibited significant time x condition interactions **(Fig. 3F & 3G, Table 1)**; homotopic connectivity between the visual seed pairs significantly increased over time in sham-sutured mice as expected due to typical development during the critical period but significantly decreased after 1 day of MD **(Table 2)**. The decrease in visual homotopic connectivity with acute MD that we observed here is consistent with depression of deprived-eye responses during OD plasticity.

**Figure 3.**
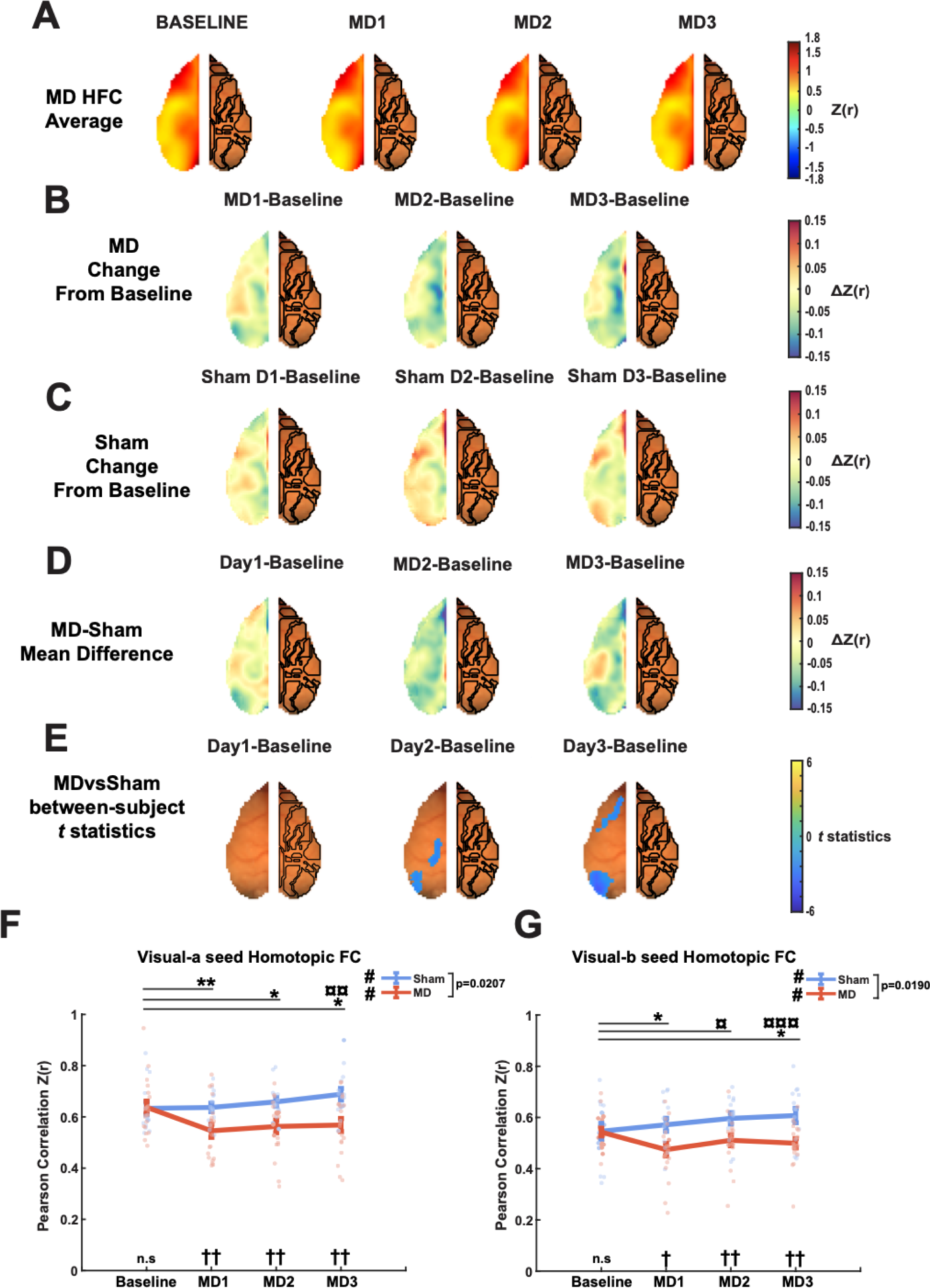
Critical period monocular deprivation (MD) alters homotopic functional connectivity patterns. **(A)** Group averages of MD homotopic FC maps across time (n=20). **(B)** MD homotopic FC plotted as the averaged change from each mouse’s baseline values. **(C)** Sham homotopic FC plotted as the averaged change from each mouse’s baseline values (n=15). **(D)** Difference maps displaying the mean difference in the changes from baseline between MD and sham mice at each MD timepoint. **(E)**. Two-sample t-tests compared the changes from baseline between MD and sham mice at each MD timepoint. **(F)** Pearson correlations between homotopic contralateral Visual-a seeds (two-way ANOVA F(3, 99) = 3.73, p = 0.021 HF-corrected). † denotes significance on post hoc two-sample t-tests comparing MD and sham mice at each timepoint (baseline: t(33) = 0.12, p = 0.90.; MD1: t(33) = −2.79, p = 0.0088; MD2: t(33) = −3.03, p = 0.0048; MD3: t(33) = −3.39, p = 0.0018). # denotes significance on rmANOVA (MD mice: F(3, 57) = 6.32, p = 0.0009; sham mice: F(3, 42) = 3.37, p = 0.027). * denotes significance on post hoc paired t-tests comparing each MD timepoint to baseline values for MD mice (MD1vsBaseline: t(19) = 3.42, p = 0.0029; MD2vsBaseline: t(19) = 2.72, p = 0.014; MD3vsBaseline: t(19) = 2.51, p = 0.021). ¤ denotes significance on post hoc paired t-tests comparing each MD timepoint to baseline values for sham mice (MD3vsBaseline: t(14) = 3.50, p = 0.0035). **(G)** Pearson correlations between homotopic contralateral visual-b seeds(two-way ANOVA F(3, 99) = 3.88, p = 0.019 HF-corrected). † denotes significance on post hoc two-sample t-tests comparing MD and sham mice at each timepoint (baseline: t(33) = −0.11, p = 0.92; MD1: t(33) = −2.71, p = 0.010; MD2: t(33) = −2.70, p = 0.011; MD3: t(33) = −3.37, p = 0.0020). # denotes significance on rmANOVA (MD mice: F(3, 57) = 4.64, p = 0.0057; sham mice: F(3, 42) = 4.66, p = 0.0067). *denotes significance on post hoc paired t-tests comparing each MD timepoint to baseline of MD mice (MD1vsBaseline: t(19) = −2.75, p = 0.013; MD2vsBaseline: t(19) = −1.35, p = 0.19; MD3vsBaseline: t(19) = −2.24, p = 0.037). ¤ denotes significance on post hoc paired t-tests comparing each MD timepoint to baseline values for sham mice (MD2vsBaseline: t(14) = 2.91, p = 0.011; MD3vsBaseline: (t14) = 4.99, p=0.0002). Error bars represent SEM here and in subsequent figures. *, ¤, † p<0.05; **, ¤¤, †† p<0.01; ¤¤¤, ††† p<0.001 here and in subsequent figures.

**Table 1.** Statistical summary of differences between MD and sham mice.

| Figures | Parameters | Data structure | Type of test | n | Statistical significance | MD effects compared to sham |
| --- | --- | --- | --- | --- | --- | --- |
| 2E | Delta GCaMP6 power maps | Normal | Two-sample t-test | n=20 MD, n=15 sham | p<0.05 for pixels displayed after cluster size-based thresholding | ↓ |
| 3E | HFC maps | Normal | Two-sample t-test | n=20 MD, n=15 sham | p<0.05 for pixels displayed after cluster size-based thresholding | ↓ |
| 3F | Visual-a seed-based FC amplitude | Normal | Two-way ANOVA | n=20 MD, n=15 sham | Time x condition interaction p=0.021 (HF-corrected) | - |
| 3F | Visual-a seed-based FC amplitude | Normal | Two-sample t-test | n=20 MD, n=15 sham | MD1 p=0.0088, MD2 p=0.0048, MD3 = 0.0018 | ↓ |
| 3G | Visual-b seed-based FC amplitude | Normal | Two-way ANOVA | n=20 MD, n=15 sham | Time x condition interaction p=0.0190 | - |
| 3G | Visual-b seed-based FC amplitude | Normal | Two-sample t-test | n=20 MD, n=15 sham | MD1 p=0.0103, MD2 p=0.0057, MD3 p=0.0020 | ↑↓ |
| 4E | Left visual-seeded FC maps | Normal | Two-sample t-test | n=20 MD, n=15 sham | p<0.05 for pixels displayed after cluster size-based thresholding | ↓ |
| 4F | Visual-parietal seed-based FC amplitude | Normal | Two-way ANOVA | n=20 MD, n=15 sham | Time x condition interaction p=0.021 (HF-corrected) | - |
| 4F | Visual-parietal seed-based FC amplitude | Normal | Two-sample t-test | n=20 MD, n=15 sham | MD1 p=0.0309, MD2 p=0.015, MD3 p=0.0019 | ↑ |
| 4G | Visual-Ss seed-based FC amplitude | Normal | Two-way ANOVA | n=20 MD, n=15 sham | Time x condition interaction p=0.014 (HF-corrected) | - |
| 4G | Visual-Ss seed-based FC amplitude | Normal | Two-sample t-test | n=20 MD, n=15 sham | MD1 p=0.021, MD2 p=0.0028, MD3 p=0.0022 | ↑ |
| S2 | Right visual-seeded FC maps | Normal | Two-sample t-test | n=20 MD, n=15 sham | p<0.05 for pixels displayed after cluster size-based thresholding | ↑↓ |

**Table 2.**
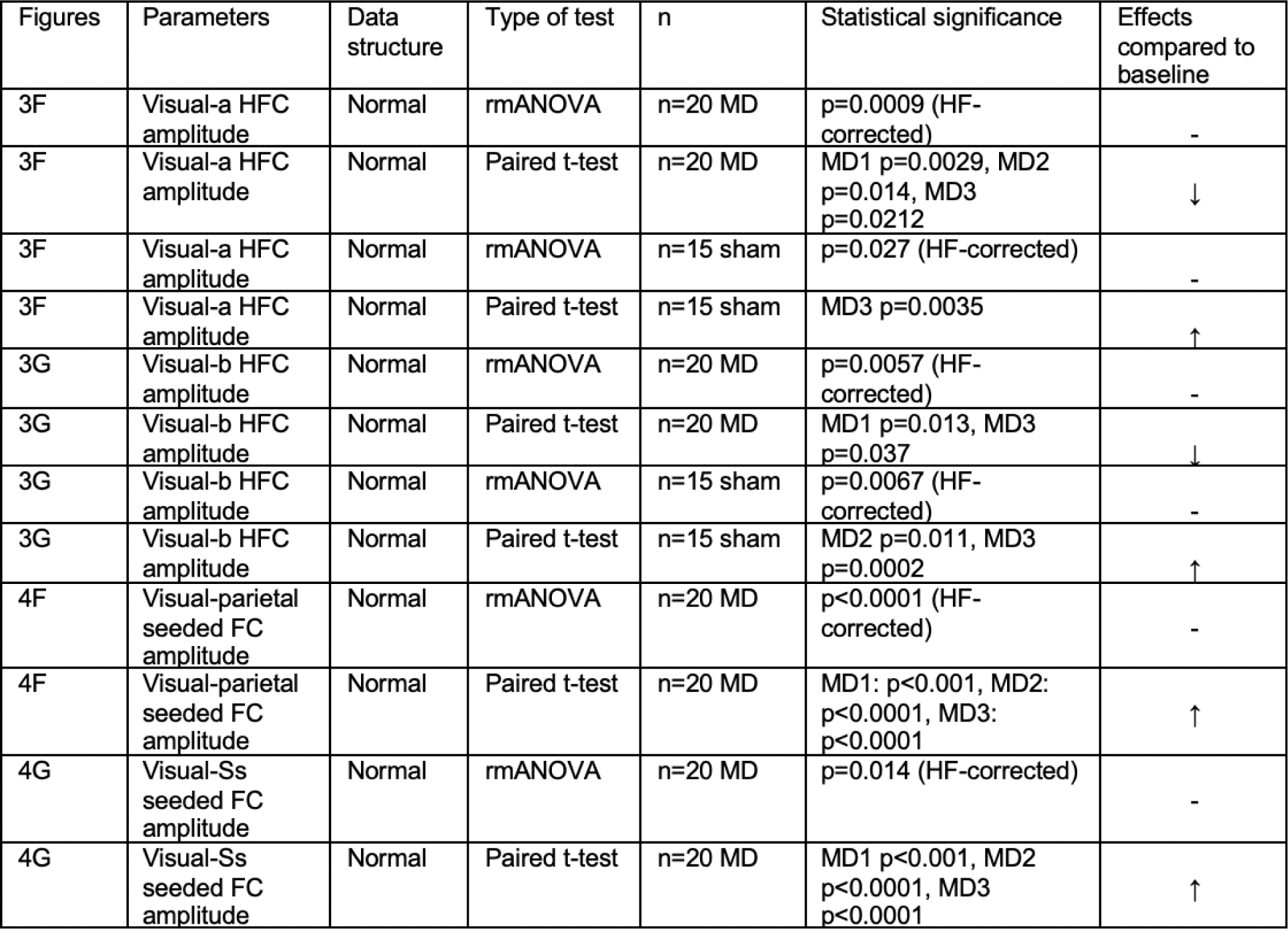
Statistical summary of differences between baseline and MD timepoints.

| Figures | Parameters | Data structure | Type of test | n | Statistical significance | Effects compared to baseline |
| --- | --- | --- | --- | --- | --- | --- |
| 3F | Visual-a HFC amplitude | Normal | rmANOVA | n=20 MD | p=0.0009 (HF-corrected) | - |
| 3F | Visual-a HFC amplitude | Normal | Paired t-test | n=20 MD | MD1 p=0.0029, MD2 p=0.014, MD3 p=0.0212 | ↓ |
| 3F | Visual-a HFC amplitude | Normal | rmANOVA | n=15 sham | p=0.027 (HF-corrected) | - |
| 3F | Visual-a HFC amplitude | Normal | Paired t-test | n=15 sham | MD3 p=0.0035 | ↑ |
| 3G | Visual-b HFC amplitude | Normal | rmANOVA | n=20 MD | p=0.0057 (HF-corrected) | - |
| 3G | Visual-b HFC amplitude | Normal | Paired t-test | n=20 MD | MD1 p=0.013, MD3 p=0.037 | ↓ |
| 3G | Visual-b HFC amplitude | Normal | rmANOVA | n=15 sham | p=0.0067 (HF-corrected) | - |
| 3G | Visual-b HFC amplitude | Normal | Paired t-test | n=15 sham | MD2 p=0.011, MD3 p=0.0002 | ↑ |
| 4F | Visual-parietal seeded FC amplitude | Normal | rmANOVA | n=20 MD | p<0.0001 (HF-corrected) | - |
| 4F | Visual-parietal seeded FC amplitude | Normal | Paired t-test | n=20 MD | MD1: p<0.001, MD2: p<0.0001, MD3: p<0.0001 | ↑ |
| 4G | Visual-Ss seeded FC amplitude | Normal | rmANOVA | n=20 MD | p=0.014 (HF-corrected) | - |
| 4G | Visual-Ss seeded FC amplitude | Normal | Paired t-test | n=20 MD | MD1 p<0.001, MD2 p<0.0001, MD3 p<0.0001 | ↑ |

### MD Results in Widespread Reorganization of Visual Cortical Connectivity

While homotopic connectivity analysis provides insight into changes in callosal connections across time, it does not assess how MD alters rsFC between visual cortex and other regions. Prior work has shown that brief (1-3d) MD shifts OD by weakening deprived eye responses without affecting the open eye responses (Frenkel & Bear, 2004; Gordon & Stryker, 1996). We therefore hypothesized that MD would specifically reorganize the connectivity between the deprived visual cortex and other non-visual regions. As a control, we expect the connectivity pattern of the non-deprived visual cortex to remain relatively unchanged.

At all timepoints, the left (deprived) visual cortex, as represented by the Visual-a seed, was highly correlated with the posterior region of the cortex and anti-correlated with the anterior region of the cortex **(Fig. 4A)**. However, when we compared each MD timepoint to baseline, we found that pixels in the parietal and somatosensory regions displayed an immediate increase in correlation to the deprived visual cortex, which persisted until MD3 **(Fig. 4B)**. To determine whether this change was due to MD or natural brain development, we repeated the analysis on sham mice. Interestingly, sham mice also displayed an increase in rsFC between the left visual cortex and the peri-parietal regions during the experiment’s time window, though to a lesser extent **(Fig. 4C)**. To directly compare MD and sham mice for the change in rsFC between left visual and peri-parietal regions, we conducted between-subject analysis on the visual rsFC maps using a two-sample t-test (**Fig. 4D)** and plotted the correlation between the left visual seed and the left parietal and somatosensory seeds **(Fig. 4E & 4F)**. Indeed, we found that the strengthening of deprived visual and peri-parietal rsFC in MD mice was significantly greater than that observed in sham mice (**Fig. 4D)**. The seed-based analyses also revealed significant time x condition interaction following a two-way rmANOVA (**Fig. 4E & 4F**, **Table 1)**; rsFC between the left visual and left parietal and somatosensory pairs significantly increased from baseline in MD mice while they did not display a significant change in sham mice **(Table 2)**. These results suggest that MD may augment certain developmental trends in this connection.

**Figure 4.**
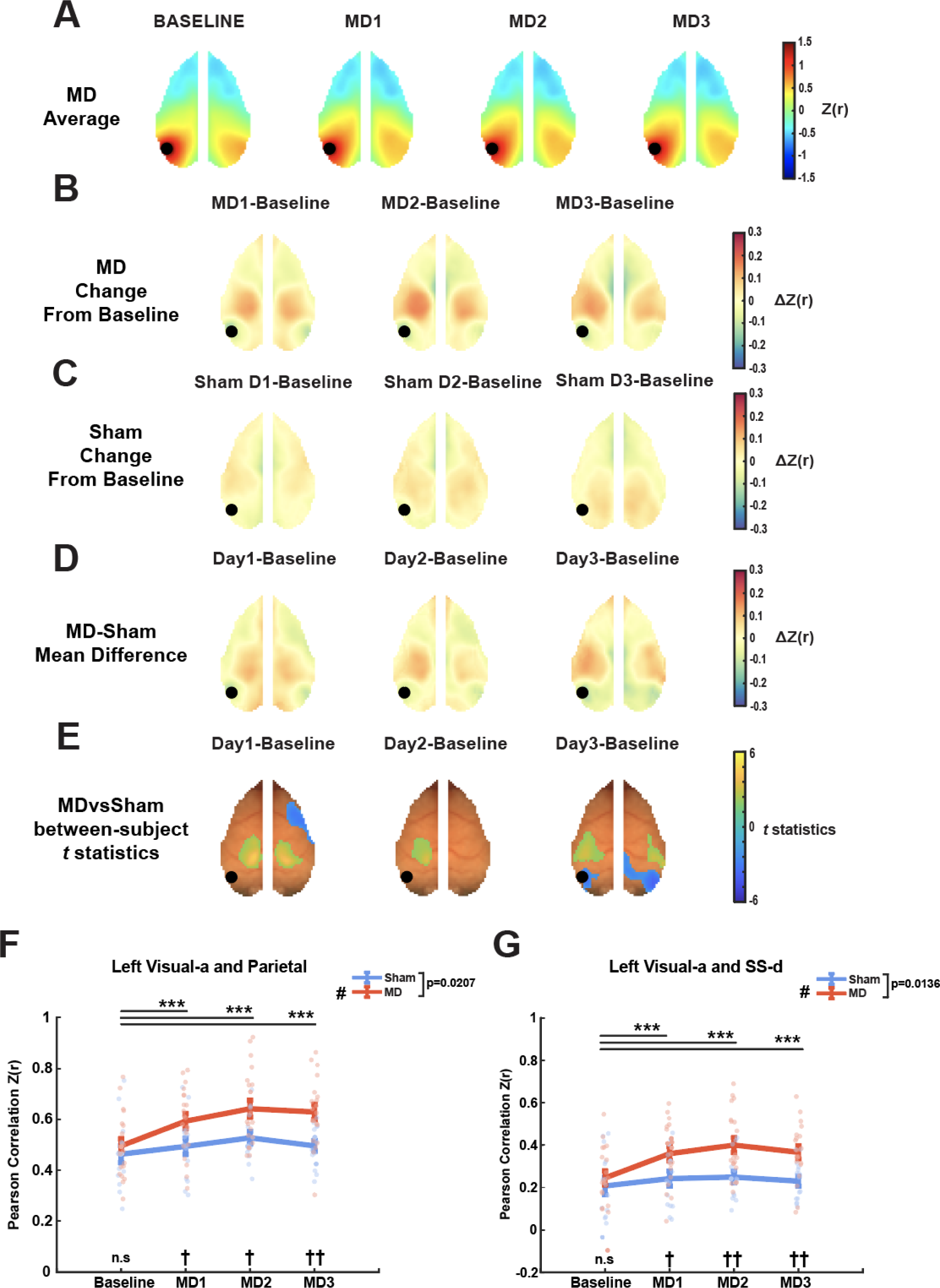
Short-term monocular deprivation alters left visual-seeded rsFC patterns. **(A)** MD average left (deprived) visual seed (represented by black dot)-based rsFC maps across time (n=20). **(B)** Pearson correlations of MD mice at each timepoint normalized to baseline values. **(C)** Pearson correlations of sham mice at each MD timepoint normalized to baseline values. **(D)** Mean difference maps illustrating the differences between normalized MD and sham mice shown in (B) and (C). **(E)** T-statistic maps showing significant clusters of pixels after two-sample t-tests comparing normalized MD and sham mice at each timepoint, thresholded using a cluster-based technique (see Methods). **(F)** Pearson correlations between left visual and parietal seeds (two-way ANOVA F(3, 99) = 3.73, p = 0.021 HF-corrected). † denotes significance on post hoc two-sample t-tests comparing MD and sham mice at each timepoint (baseline: t(33) = 0.74, p = 0.47; MD1: t(33) = 2.25, p = 0.031; MD2: t(33) = 2.58, p = 0.015; MD3: t(33)= 3.38, p = 0.0019). # denotes significance on rmANOVA (MD mice: F(3, 57) = 16.354, p < 0.0001). *denotes significance on post hoc paired t-tests comparing each MD timepoint to baseline of MD mice (MD1vsBaseline: t(19) = 3.96, p = 8.4e-4; MD2vsBaseline: t(19) = 4.9005, p = 9.9e-5; MD3vsBaseline: t(19) = 5.86, p = 1.22e-5). **(G)** Pearson correlations between left visual and somatosensory-d seeds (two-way ANOVA, F(3, 99) = 3.92, p = 0.014 HF-corrected). † denotes significance on post hoc two-sample t-tests comparing MD and sham mice at each timepoint (baseline: t(33) = 0.73, p = 0.47; MD1: t(33) = 2.42, p = 0.021; MD2: t(33) = 3.23, p = 0.0028; MD3: t(33) = 3.32, p = 0.0022). # denotes significance on rmANOVA (MD mice: F(3, 57) = 14.00, p < 0.0001). *denotes significance on post hoc paired t-tests comparing each MD timepoint to baseline of MD mice (MD1vsBaseline: t(19) = 4.92, p = 9.4e-4; MD2vsBaseline: t(19) = 4.63, p = 1.8e-5; MD3vsBaseline: t(19) = 4.49, p = 2.5e-5).

To confirm that these changes are specific to the deprived visual cortex, we examined changes in the right visual-seeded FC maps **(Fig. S1)**. Following MD, the non-deprived visual cortex exhibited a significant decrease in its correlation to the deprived (contralateral) visual cortex and a transient but significant increase in its correlation to the ipsilateral motor region **(Fig. S1B & S1E)**. However, there were no significant changes in rsFC between the non-deprived visual cortex and the peri-parietal region, which suggests that the loss of input induces changes in connectivity specific to the deprived visual cortex.

Taken together, these data indicate that critical period MD has transcallosal impacts on rsFC, rapidly strengthening rsFC between the deprived visual cortex and peri-parietal regions on both hemispheres and weakening rsFC between the visual cortices as well as between the deprived visual cortex and the contralateral motor region. The reduced connectivity between the visual regions may be remapped to strengthen connectivity between visual and other non-sensory/associative regions.

## Discussion

The responsiveness of visual cortex neurons is modulated by several distinct mechanisms during MD, but the effects of MD on neocortical structures beyond visual cortices are not well-documented. Therefore, we sought to determine whether MD has circuit-level impacts across the cortex. Using a classic MD and calcium-based optical imaging paradigm, we showed that critical period MD impacts rsFC between several cortical regions. Because MD induces a discordance in inputs received by the two hemispheres, changes in callosal connections likely contribute to alterations in correlated spontaneous activity in MD mice. Prior work has shown that MD activates interhemispheric inhibition where inputs from the corpus callosum suppress closed eye responses thereby shifting OD toward the open eye (Pietrasanta et al., 2014; Restani et al., 2009), and enhanced inhibitory activity has been thought to decrease the strength of rsFC between homotopic regions (Antonenko et al., 2017; Stagg et al., 2014). Here, we have shown that a lower visual homotopic functional connectivity occurred as early as two days after MD. MD-induced activation of interhemispheric inhibition is consistent with this decrease in spontaneous activity in the visual cortices.

Our data suggest that the correlation between the deprived visual cortex and the peri-parietal regions in both hemispheres increased immediately following MD. The parietal cortex is a multi-modal association region implicated in visuo-spatial perception, spatial navigation, and movement control (Krumin et al., 2018; Lyamzin & Benucci, 2019; Oh et al., 2021). Disruptions of parietal cortex activity impair visually-guided decision making (Licata et al., 2017). The higher rsFC between the deprived visual cortex and peri-parietal regions may reflect changes in how sensory information is associated and integrated for guiding behavior. Indeed, the distinct effects seen in left versus right visual rsFC suggest that the correlation of calcium activity matches the imbalance of sensory input. It’s worth noting that sham mice showed a similar trend where rsFC between the visual cortex and peri-parietal region increased, though to a lesser extent. Since P24-P27 is within the mouse visual critical period (Reh et al., 2020), the baseline increase in sham animals suggests that as sensory integration strengthens, connectivity between the visual cortex and those regions responsible for integration and non-visual sensory functions increases. Our deprivation results further suggest that instead of working against this developmental trend, MD sped up this process. In marked contrast, maps from the seed in the non-deprived visual cortex showed no signs of increasing connectivity to either the left or right parietal regions **(Fig. S1)**. For further context on parietal connectivity, in our previous longitudinal study of brain development across P15 through P60 (Rahn et al., 2021), we observed a slower continuing increase in contralateral connectiviy between left-right parietal regions, where in contrast almost all other contralateral connections which peaked at around P21 or P35. Together these results suggest the parietal regions play an integral yet complex role in the FC developmental trajectories.

Additionally, we found MD resulted in greater anti-correlation between the deprived visual cortex and the contralateral motor cortex. The enhanced visual-motor anticorrelation has previously been observed after 14 days of MD (Kraft et al., 2017). Although consistent with the prior prediction that the enhanced visual-motor anti-correlation may be the result of adapting behavior to an imbalanced visual experience (Kraft et al., 2017), our findings here may also suggest the rewiring process occurs much earlier than previously reported.

Finally, beyond connections from visual seeds to the rest of the brain, we also examined if any other seed pairs changed connection strength across the whole cortex by calculating the correlation between both homotopic and non-homotopic seed pairs and their changes across time **(Fig. S2)**. While there were some changes seen in individual connections (including the expected changes in visual connections detected above), no additional significant differences in connectivity were detected after applying statistical correction for the number of hypothesis tests (n=325) this brain-wide analysis entailed. Thus any such indirect changes would require additional power to identify.

Previous experiments have demonstrated that while most visual cortex neurons have stable firing properties across circadian light-dark (L-D) transitions and across sleep and wake states, disruptions of the 12:12 L-D cycle can lead to changes in the activity of some visual neuronal populations (Cary & Turrigiano, 2021; Torrado Pacheco et al., 2019). If collecting imaging data across L-D cycles and across behavioral states chronically were to become feasible, such as through the use of miniscopes, future studies could investigate the presence (or absence) of environmental stimuli and sleep and wake cycles on the reorganization of brain connectivity patterns during MD. We have collected pilot data where mice were exposed to a 24:0 L-D cycle and found the increased rsFC between the deprived visual cortex and peri-parietal regions occurred faster and to a higher extent than those exposed to 12:12 L-D cycles (data not shown).

Overall, the results of this study suggest that sensory deprivation during critical developmental windows can reprogram the cortex and alter FC between disparate regions. While we focus on the visual network, future studies could test whether loss of input to other sensory systems, such as the whisker barrels, similarly redirects their functional connectivity to associative cortices. Finally, as with FC studies in general, it is also of interest to understand if these changes in correlated activity represent alterations in direct anatomical connectivity. The studies here motivate further investigation into potential remapping of connectivity in distal cortico-cortical connections in response to sensory deprivation during critical windows of development.

## Author Contributions

S.C., R.M.R., K.B.H., J.D.D., and J.P.C. designed research; S.C., A.R.B., S.H.B. performed research; S.C., R.M.R., J.A.P.C, and J.P.C. contributed analytical tools; S.C. and J.A.P.C. analyzed the data; S.C., R.M.R., A.R.B., S.H.B., J.A.P.C., K.B.H., J.D.D., and J.P.C. wrote the paper.

## Acknowledgements

This work was supported by the National Institute of Health (R01MH124808, R01MH107515 to JDD and R01NS099429 to JPC) and Washington University Intellectual and Developmental Disability Research Center (P50HD103525 to JDD). We thank Adam Q. Bauer for helpful discussions and advice.

## Supplementary Information

**Figure S1.**
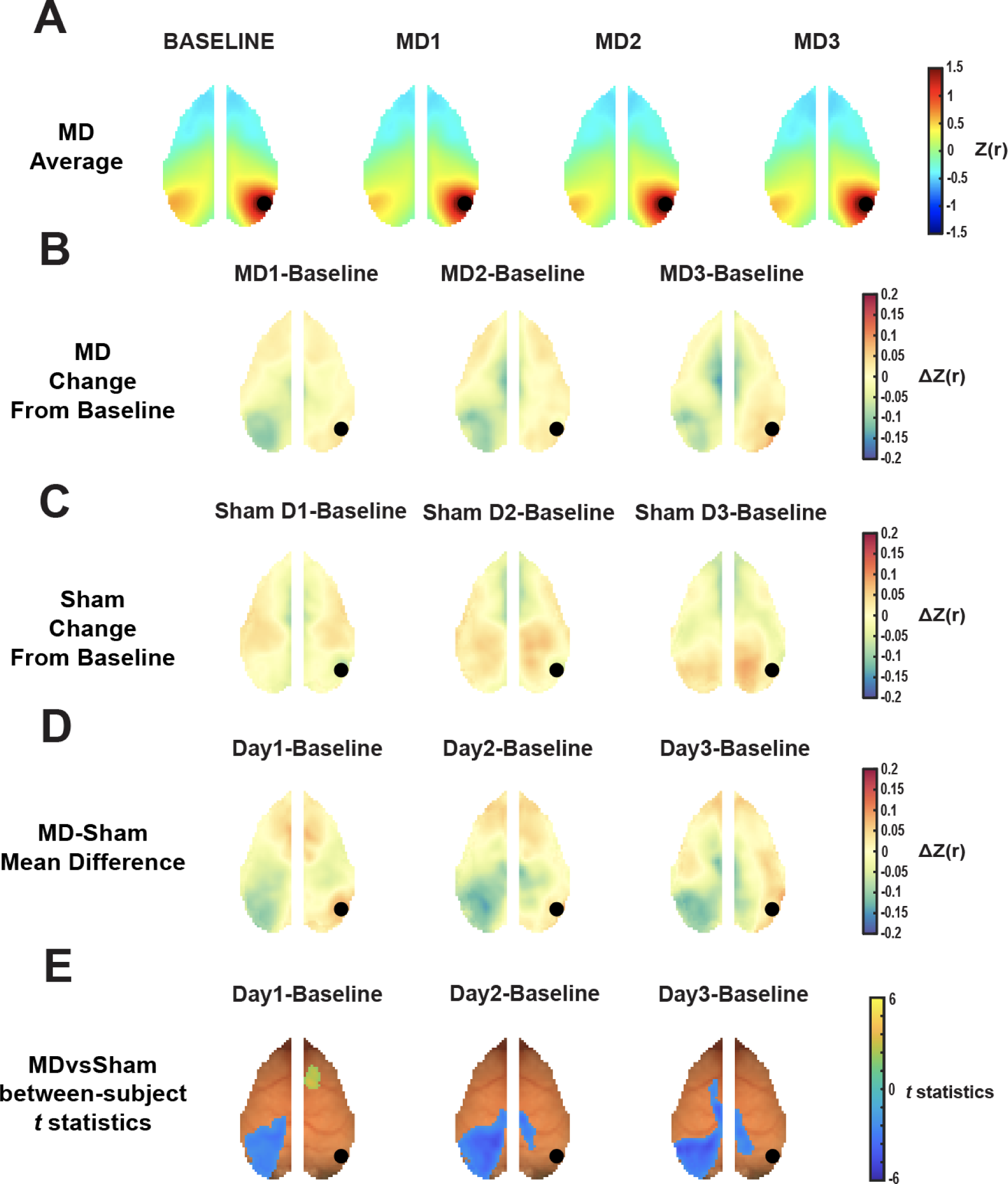
The non-deprived visual cortex exhibited decreased correlation to the deprived visual cortex. **(A)** MD average right (non-deprived) visual seed (represented by black dot)-based rsFC maps across time (n=20). **(B)** Pearson correlations of MD mice at each timepoint normalized to baseline values. **(C)** Pearson correlations of sham mice at each MD timepoint normalized to baseline values (n=15). **(D)** Mean difference maps illustrating the differences between normalized MD and sham mice shown in (B) and (C). **(E)** T-statistic maps showing significant clusters of pixels after two-sample t-tests comparing normalized MD and sham mice at each timepoint, thresholded using a cluster-based technique.

**Figure S2.**
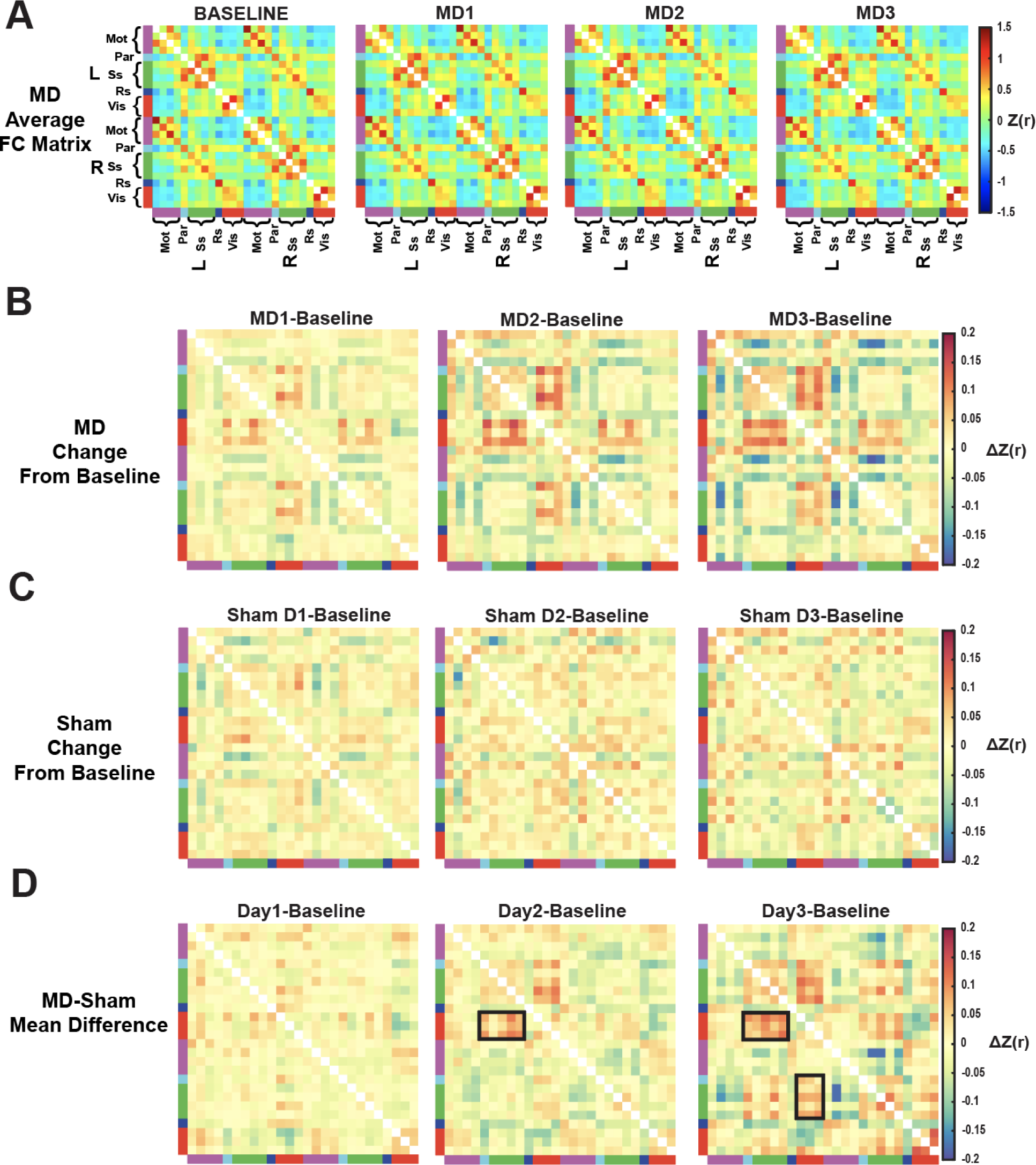
rsFC Matrices demonstrated an increase in correlation between visual and sensory networks. **(A)** MD average rsFC matrices representing Pearson correlations between network seed pairs (n=20). **(B)** Pearson correlations normalized to baseline values for MD mice. **(C)** Pearson correlations normalized to baseline values for sham mice (n=15). **(D)** Mean difference between normalized MD and sham groups. Boxes highlight increased correlation between visual and parietal and somatosensory networks.

